# Complementary methods for the study of interactions between eosinophils and cancer cells

**DOI:** 10.1101/2024.04.25.590232

**Authors:** Caterina Antonucci, Adriana Rosa Gambardella, Valentina Tirelli, Fabrizio Mattei, Giovanna Schiavoni

**Affiliations:** Department of Oncology and Molecular Medicine, Istituto Superiore di Sanità, Viale Regina Elena, 299, 00161, Rome, Italy; Core Facilities, Istituto Superiore di Sanità, Viale Regina Elena, 299, 00161, Rome, Italy

## Abstract

Eosinophils are a rare immune cell subset with important roles in Th2 immunity and, recently, in cancer. Interleukin IL-33 (IL-33) is well recognized for its important roles in the activation of eosinophils in Th2 immunity. On the other hand, IL-33 has been recently discovered to play central roles in cancer, in particular by activating eosinophils and increase their degranulation consequent to an intrinsic tumor cell killing function. We propose a dual approach methodology to extrapolate functional interactions of eosinophils with tumor cells, as a result of eosinophil stimulation. Human eosinophils (Eos) isolated from the blood of healthy donors by dextran sedimentation followed by magnetic sorting are exposed to IL-33 (Eos33) or IL-5 (Eos5, control) for 18 h. These pre-conditioned cells are then co-cultured with A375P melanoma cells to monitor cell-cell interactions. Acoustic focusing flow cytometry analysis is employed to evaluate the presence of Eos-tumor cell conjugates after 1h incubation of human eosinophils and A375P melanoma cells. Moreover, a 24 h time-lapse video recording approach is employed to obtain single cell tracking Eos profiles. This allows to quantitatively determine the interaction extent of Eos33, as opposed to Eos5 (control condition), with tumor cells. In conclusion, our protocols easily and quickly allow the extrapolation of relevant kinematic and biologically relevant parameters for tumor reactive eosinophils. Furthermore, these methods are adaptable to various models with other types of immune cell subsets and cancer cells and can be implemented on different video microscopy platforms and advanced flow cytometry systems.

## 1. Introduction

Eosinophils (Eos) are a rare population of granulocytes that represent 1-3% of total blood leukocytes under physiological conditions. Eos play a prominent roles in allergic and parasitic infection but are also involved in angiogenesis and tumor immunity (Rosenberg et al., 2013). Eos release a plethora of soluble mediators, among which Eosinophil Cationic Protein (ECP), Eosinophil Peroxide (EPX) and Eosinophil-derived Neurotoxin (EDN), cytokines, chemokines, growth and pro-angiogenic factors and variably affect cancer progression (Nissim Ben Efraim & Levi-Schaffer, 2014; Varricchi, Galdiero, et al., 2018; Varricchi, Loffredo, et al., 2018). Human Eos originate from the bone marrow, where they differentiate from progenitor cells to mature Eos through various stimuli. The process of proliferation and differentiation is first guided by the GATA-binding Factor 1 transcription factor (GATA-1). At this point, IL-5 together with IL-3 and granulocyte-macrophage colony-stimulating factor (GM-CSF) sustain the development and maturation of the Eos inside the bone marrow microenvironment (Varricchi, Galdiero, et al., 2018). Once Eos become mature and express the IL-5Ra, CCR3, EMR1, CRTH2, and Siglec-8 receptors, they leave bone marrow tissue and mobilize to the bloodstream from where they can reach target tissue in response to danger stimuli.

Increasing evidence indicates that the alarmin IL-33 is required for both basal homeostasis (Johnston et al., 2016) and activation of eosinophils (Cherry et al., 2008). This cytokine regulates Eos at multiple levels, during their maturation and activation inducing up-regulation of adhesion molecules (i.e., CD11a and CD11b) and the activation marker CD69, pro-inflammatory cytokines and chemokines, superoxide anion production and degranulation (Andreone et al., 2019; Cherry et al., 2008; Hashiguchi et al., 2015). In melanoma models, IL-33 promotes anti-tumor immunity through recruitment and activation of Eos, which induce contact-dependent degranulation and tumor cell killing (Andreone et al., 2020; Andreone et al., 2019; Lucarini et al., 2017). In the tumor microenvironment, Eos have been detected interacting with cancer cells in several human malignancies (Caruso et al., 2011; Ghaffari & Rezaei, 2023; Varricchi, Galdiero, et al., 2018). In this respect, direct contact of Eos with a target tumor cells is a pre-requisite for effector function of these cells, such as cytotoxic degranulation (Andreone et al., 2019; Gatault et al., 2015) and trogocytosis (Mattei et al., 2022).

Based on these premises, we present a protocol to investigate the direct interactions of Eos with tumor cells through two different complimentary methods. Eosinophils isolated from the peripheral blood are stimulated with the conventional IL-5 exposure (Eos5, control condition) or with IL-33 (Eos33), and then co-cultured with A375P human melanoma cells. Acoustic focusing flow cytometry analysis (Lugo-Gavidia et al., 2021) provides a precise quantification and imaging of the conjugates between labelled A375P melanoma cells and Eos33 as opposed to Eos5. In parallel, time lapse video recording enables the visual and quantitative observation in real time of the interactions between Eos5 or Eos33 with A375 cells by extrapolation of meaningful tracking profiles of eosinophil trajectories and associated parameters. Conclusively, these methods could constitute an alternative but reliable approach to evaluate anti-cancer activities of eosinophils that can easily be adapted to other immune effector cells interacting with tumor cells.

## 2. Materials

### 2.1 Common disposable

- 50 mL Polystyrene conical-bottom centrifuge tubes (#ET5050B, Euroclone)
- 15 mL Polystyrene conical-bottom centrifuge tubes (#ET5015B, Euroclone)
- FACS tubes (e.g., 5 mL Polystyrene round-bottom tubes with cap, #352054, Corning)
- Sterile culture flasks with filter, 25 cm^2^ (#ET7026, Euroclone)
- Sterile culture flasks with filter, 75 cm^2^ (#ET7076, Euroclone)
- Cell count slides with counting grids (#87144, KOVA International)
- LS Columns (#130-042-401, Milenyi Biotec)

### 2.2 Cells and Reagents

- A375P melanoma cell line (#CRL-3224, ATCC)
- Buffy coats from human healthy donor peripheral blood (Prot. n. OO-ISS 02/10/2019 0029604 approval by the Ethics Committee of the Istituto Superiore di Sanità, Rome, Italy)
- IL-5 (#GFH-191-100, Cell Guidance Systems)
- IL-33 (#GFH145-10, Cell Guidance Systems)
- RPMI 1640 (#ECB9006L, EuroClone)
- DMEM High Glucose (#ECB7501L, Euroclone)
- FBS (Fetal Bovine Serum South America origin EU Approved, #ECS5000L, EuroClone)
- Antibiotic-Antimycotic Solution 100X (10000 U/mL Penicillin G, 10,000 μg/mL Streptomycin, 25 μg/mL Amphotericin B; #SV3007901, Euroclone)
- L-Glutamine 100X (#ECB3003D, Euroclone)
- HEPES Buffer 1M (#ECM0180D, Euroclone)
- NEAA (Non Essential AmminoAcids 100X Solution; #ECB3054D, Euroclone)
- Sodium pyruvate 100 mM (#ECM0542D, Euroclone)
- Dextran 70 (#A1847, Biochemica)
- Dextran/NaCl: 0.6 M Dextran, 0.154 M of NaCl (see *Note 1*)
- ACK, Ammonium-chloride potassium red blood cell lysis buffer (see *Note 2*)
- Lymphosep, Lymphocyte Separation Media (#AU-L0560500, Biowest)
- PBS 1X, Phosphate-Buffered Saline (#ECB4004L, Euroclone)
- Eosinophil medium: RPMI 1640 supplemented with 20% FBS, 1% glutamine, 1% Antibiotic-Antimycotic, 1% HEPES, 1% NEAA, 1% Sodium pyruvate
- DMEM culture medium: DMEM High Glucose supplemented with 10% FBS, 1% glutamine, 1% Antibiotic-Antimycotic
- RPMI co-culture medium: RPMI 1640 supplemented with 10% FBS, 1% glutamine, 1% Antibiotic-Antimycotic
- Macs Buffer: 1% FBS, 2 mM EDTA in PBS 1X
- Facs Buffer: 1 g/L NaN_3_, 2% FBS in PBS 1X
- Trypan Blue 0.4% solution (dilute 1:8 in PBS 1X; #17492E, Lonza)
- Eosinophil Isolation Kit, human (Miltenyi Biotec, #130-092-010)
- Human Siglec-8 antibody, conjugated to PE-Vio 770 (#130-105-524, Miltenyi Biotec)
- PKH26 Red fluorescent Cell Linker (#MINI26, Sigma-Aldrich)

### 2.3 Equipment

- MidiMACS™ Separator (#130-042-302, Miltenyi Biotec)
- MACS® MultiStand (#130-042-303, Miltenyi Biotec)
- Class II Laboratory biosafety cabinet VBH 75 MP (#8071, Steril)
- SL 1R Plus MD centrifuge with full equipment (#17190589, Thermo Scientific)
- Centrifuge 5471R with full equipment (#EP022620701, Eppendorf)
- Attune™ CytPix Acoustic Focusing Flow Cytometer (#49434, Thermo Scientific)
- Telaval 31 Optical inverted microscope (#3514, Zeiss)
- Forma Direct Heat CO_2_ Incubator (#310TS, Thermo Scientific)
- JuLI Smart Fluorescent videomicroscopy system (#08011, BulldogBio)

### 2.4 Software

- ImageJ/Fiji v.1.54f with Java v1.8.0_172 (National Institutes of Health, USA; https://imagej.net/ij/)
- ImageJ plugin TrackMate (https://imagej.net/TrackMate)
- Java object-oriented programming language (Oracle; https://www.java.com)
- Attune Cytometric Software v6.1.0 (Thermo Fisher Scientific)
- Prism GraphPad v.6 (https://www.graphpad.com/)

## 3. Methods

### 3.1 Purification of human Eos from peripheral blood

1. Prepare 50 mL tubes each with 6.25 mL pre-warmed Dextran/NaCl plus 2.5 mL EDTA 0.1 M (pH 7.4) (see *Note 3*).
2. Add 10 mL of peripheral blood buffy coat to each 50 mL tube and mix by gently reverse the tube 3/4 times
3. Incubate the tubes at room temperature (RT) to allow leukocyte sedimentation into the Dextran/NaCl gradient.
4. Start to collect the upper leukocyte portion (white cells) after 15 min and up until 30-35 min and transfer in a new 50 mL tube containing 10 mL of PBS 1X (see *Note 4*).
5. Bring up to 50 mL volume with PBS 1X
6. Centrifuge the tubes at 1500 rpm, 6 min at RT.
7. Discard the supernatant and resuspend the leukocyte pellet in 40 mL PBS 1X (see *Note 5*).
8. Centrifuge at 1200 rpm for 8 min at RT (see *Note 6*).
9. Discard the supernatant and resuspend the leukocyte pellet in 5 mL PBS 1X at RT (see *Notes 6 and 7*).
10. Prepare a 15 mL tube containing 5 mL Lymphosep Lymphocyte Separation Media.
11. Carefully stratify the leukocyte suspension onto the 5 mL Lymphosep at RT (see *Note 8*).
12. Centrifuge at 1800 rpm, 20 min without brake at RT (see *Note 9*).
13. Carefully aspirate the supernatant by using a serological pipette and discard (see *Note 10*).
14. Collect the pellet that contains granulocytes with 5 mL PBS 1X.
15. Transfer the resuspended pellet in a new 50 mL tube containing PBS 1X.
16. Bring up to 50 mL PBS 1X and centrifuge at 1200 rpm for 8 min at RT.
17. Discard the supernatant and repeat the wash in the previous step (see *Note 11*).
18. Discard the supernatant, gently resuspend the pellet in 5 mL ACK and incubate for 5 min at R.T. During this incubation mix the tube at least twice by gentle agitation (see *Note 12*).
19. Stop the ACK reaction by adding a two-fold amount of ice cold Macs Buffer (see *Note 13*).
20. Centrifuge at 1300 rpm for 5 minutes at 4°C.
21. Discard the supernatant and resuspend the pellet of granulocytes in ice cold Macs Buffer. Keep cell suspension on ice (see *Notes 13 and 14*).
22. Count granulocytes by Trypan Blue exclusion (see *Note 15*).
23. Use the Miltenyi Human Eosinophils Isolation Kit for immunomagnetic separation of untouched human Eos from the granulocyte suspension as per manufacturer’s instructions (see *Note 16*).
24. Collect the negative fraction of cells that pass through the LS column (flow-through) that constitute the isolated human eosinophil population.
25. Pellet the flow-through cells by centrifuging 1600 rpm, 5 minutes, 4°C.
26. Discard the supernatant and resuspend in appropriate volume of Eosinophil culture medium and count cells (see *Note 15*).

### 3.2 Stimulation of Eos with IL-5 or IL-33

1. Adjust concentration of Eos obtained (Section 3.1, step 26) to 10^6^ cells/mL.
2. Seed 1-2 × 10^6^ Eos per well in 24 well culture plates for a total of at least two wells (see *Note 17*).
3. Stimulate Eos with either 10 ng/mL of recombinant human IL-5 (Eos5) or 50 ng/mL of recombinant human IL-33 (Eos33) by adding the cytokines to the wells.
4. Incubate 18 h at 37°C and 5% CO_2_.
5. Collect the Eos (thereafter Eos5 and Eos33) and centrifuge at 1600 rpm, 5 min, 4°C.
6. Resuspend the pellets in Eos culture medium and count cells by Trypan blue exclusion (see *Note 15*).
7. Adjust to the desired volume and store at 4°C until use.

### 3.3 Tumor cell culture

1. A375P cells are routinely cultured in DMEM culture medium.
2. When cells achieve 70–80% confluence, exhausted culture medium is discarded by aspiration, and cells are gently washed with 0.5 mL pre-warmed PBS 1X
3. PBS 1X is discarded by aspiration and cells are detached by incubation in 2 mL 0.05% Trypsin-EDTA solution for 1–3min at 37°C
4. Detached cells are harvested and centrifuged at 1500 rpm for 5 min, followed by aspiration of the supernatant
5. Cells are resuspended in complete culture medium for further use.

### 3.4 PKH26 Red fluorescent Cell Linker staining

1. For conjugate evaluation by flow cytometry, A375P cells are labelled with the PKH26 fluorescent cell tracker. To this purpose, cells are resuspended in PBS 1X at the concentration of 10^7^ cells/mL (eg. 0.5 × 10^6^ cells in 500 μl).
2. Prepare an equal volume (eg. 500 μL) of 2X PKH26 Red fluorescent Cell Linker Solution in the dark at RT (see *Note 18*).
3. Add the 2X PKH26 Red fluorescent Cell Linker solution to the A375P cell suspension, mix thoroughly and incubate for 5 min in the dark at RT (see *Note 19*).
4. Add 1 ml FBS to stop the labeling reaction and incubate for 1 min in the dark at RT.
5. Pellet cells by centrifuging 1500 rpm for 10 min at 4°C and discard the supernatant.
6. Wash the A375P cells with 5 mL of DMEM culture medium and pellet them by centrifuging at 1500 rpm for 5 min at 4°C.
7. Discard the supernatant, resuspend the pellet in DMEM culture medium and count cells (see *Note 15*).

### 3.5 Co-culture of Eosinophils and A375 cells for conjugate evaluation

1. In 5 ml tubes (two tubes needed) seed PKH26-labelled A375P cells (Section 3.4, step 7) 10^5^/tube resuspended in 100 μl of RPMI co-culture medium.
2. Resuspend Eos5 or Eos33 in RPMI co-culture medium at 10^7^ cells/mL.
3. Add 100 μL of either Eos5 or Eos33 (Section 3.2, step 7) corresponding to 10^6^ cells/tube to reach a 10:1 (Eos:A375P) ratio.
4. Gently mix the tubes and place in the incubator at 37°C with 5% CO_2_ for 1h leaving the lid loose (see *Note 20*).

### 3.6 Siglec-8 labelling of Eos5/PKH26^+^A375P and Eos33/PKH26^+^A375P co-cultures

1. At the end of the 1h incubation (Section 3.5, step 4), co-cultures are centrifuged at 1600 rpm for 5 min at 4°C (see *Note 21*).
2. The supernatant is discarded and the cell pellet is resuspended in 1-2 mL ice cold FACS Buffer (see *Note 22*).
3. Centrifuge cells at 1300 rpm for 5 min at 4°C.
4. Discard the supernatant, gently resuspend the pellet and add 2 μL PE-Vio 770 Siglec-8 antibody directly to the pellet (see *Note 23*).
5. Incubate cells for 15’ at 4°C in the dark.
6. Wash cells with 1-2 mL FACS Buffer and centrifuge at 1300 rpm for 5 min at 4°C.
7. Discard the supernatant and resuspend the cell pellet in 400 μL FACS Buffer (see *Note 22*).
8. The two Siglec-8^+^Eos5/PKH26^+^A375P and Siglec-8^+^Eos33/PKH26^+^A375P co-culture conditions are then ready to be acquired by the Attune CytPix Acoustic Focusing flow cytometer for Eos/A375P conjugates detection and imaging (Section 3.7).

### 3.7 Acquisition of cells with the Attune CytPix Acoustic Focused Flow Cytometer

1. Flow cytometry acquisition is performed on an Attune CytPix Flow Cytometer equipped with 488 nm (blue), 640 nm (red), 405 nm (violet), and 561 nm (yellow) lasers and a high-speed brightfield camera that captures and stores images of detected events as samples are acquired at a rate of up to 6,000 images/second (see *Note 24*).
2. To acquire data, the following channels are used: SSC, FSC, YL1 (561 nm laser, 780/60 nm filter) and YL4 (561 nm laser, 780/60 nm filter)
3. Laser voltage (gain) and compensation are set using unstained cells and cells stained with each fluorochrome separately.
4. At least 10,000 events with normal SSC and FSC are acquired per sample, ideally at a flow rate < 500 events/second.

### 3.8 Analysis of Eos/A375P conjugates with the Attune Cytometric Software

1. In the Attune Cytometric software generate FSC/SSC, FSCA/FSCH and Siglec-8/PKH26 dot plot masks.
2. The different samples are analyzed with the same parameters to identify the presence of conjugates A375P/Eos5 and A375P/Eos33 in the samples.
3. The full gating strategy to discriminate the conjugates by the single cells begins with the selection of viable cells based on FSC/SSC morphological parameters (gate object “morpho”).
4. Select events with higher cell size and granularity based on FSCA/FSCH plot (gate object “large”).
5. Identify double positive PKH26Red^+^Siglec8^+^ cells from the Siglec-8/PKH26 plot (gate “conjugates”) (see *Note 25*).
6. From the gate “conjugates” generate images for each event. Actual eosinophil-A375P conjugate are visualized and selected based on morphometry and counted manually. Images portraying Eos5 or Eos33 attached to A375P are selected based on morphometry, while false positives (i.e., dead cells, unattached eosinophil-tumor cell, or tumor cell doublets) are discarded (see Figure 1).

**Figure 1.**
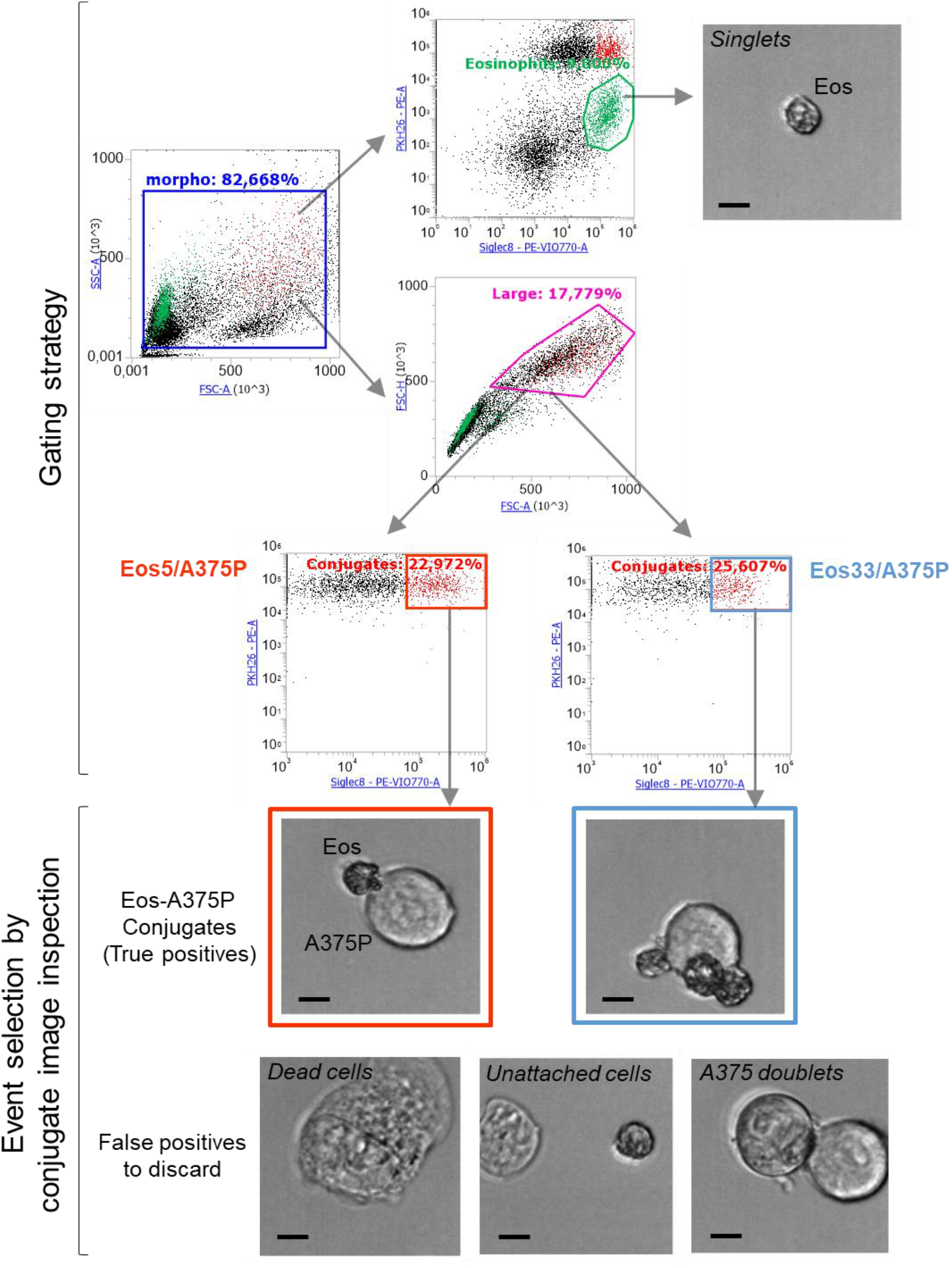
Analysis of Eos/A375P conjugates after 1 h co-culture by Attune CytPix Acoustic Focusing Flow Cytometer. Representation of the gating strategy employed to Eos5/A375P or Eos33/A375P conjugates. Morphological fit cells are identified in the FSC/SSC plot (“morpho” gate). The “morpho” gated cells plotted in PKH26/Siglec-8-PE-Vio770 allow to distinguish single eosinophils (PKH26^-^Siglec-8^+^) and PKH26+ tumor cells. To enrich the population containing cell conjugate plot FSCA/FSCH and select large cells (“large” gate) population include PKH26^+^Siglec-8 ^+^ doublets (“conjugates”). Conjugates in the two Eos5/A375P and Eos33/A375P experimental conditions (Red and blue rectangular gates) are delineated by some representative images acquired by automatic focused visible light microphotograph during the clustering process of eosinophil singlets (i.e., Eos single cells in green gate), doublets (True positives with effective Eos-A375P contact) and the indicated false positives. Image inspection allows to reveal an estimate of the false positive rate and to discard them. Small, phase contrast-dark cells in the images represent granule-containing eosinophils, whereas larger cells are the A375P tumor cells. Scalebar, 10 micrometers.
7. Count the eosinophil-A375 conjugates formed by one tumor cells and one or more Eos5/ Eos33 and extrapolate the percentages on total eosinophils. In our specific example, conjugates containing two or more Eos5 and A375P represent the 23,26 % and conjugate with two or more Eos33 and tumor cell represent the 31,5% (see Figure 1 and *Note 26*).

### 3.9 Co-culture of Eosinophils and A375 cells for time-lapse video recording

1. One day before the time lapse start, place the two Juli Smart microscopes into the CO_2_ incubator (Figure 2A) (see *Note 27*)

**Figure 2.**
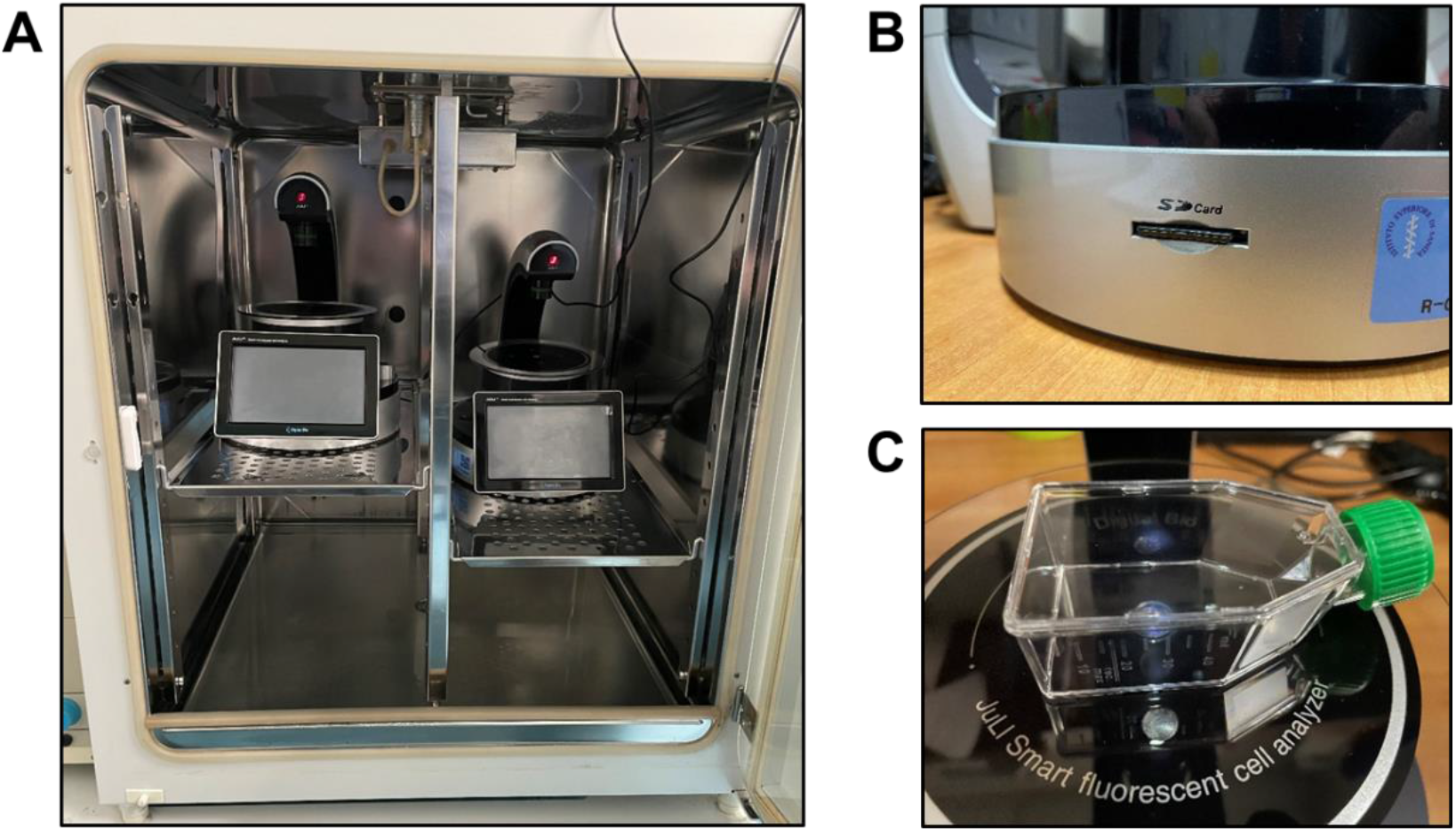
Use of the Juli Videomicroscopy Systems for the generation of time lapse video. (A) The two Juli Smart Fluorescent Analyzers positioned at the inner of the CO_2_ incubator before starting the time lapse. Black electric wires exiting from the incubator provide power current to microscopes. (B) Detail of the SD card slot on a Juli Smart Fluorescent Analyzer, with a SD card inserted and ready to acquire images. (C) Correct positioning of a 25 cm^2^ cell culture flask on the Juli stage.
2. In the same day, seed 10^5^ A375P cells (see section 3.3) in 25 cm^2^ Flasks (two flasks needed) and incubate at 37°C 5% CO_2_ for 18-20 h (overnight).
3. The following day, 3 × 10^5^ Eos5 or Eos33 (see Section 3.2) are alternatively added to the 25 cm^2^ Flask containing A375P cells in order to achieve a 3:1 (Eos:A375P) ratio (see *Notes 28 and 29*).
4. Insert the SD card (see *Note 30*) in the appropriate outlet of the microscope (Figure 2B) and turn the instrument on.
5. Place the two flasks containing Eos5/A375P and Eos33/Ar35P co-cultures Add Eos on the stage of the microscope, as depicted in Figure 2C.
6. The time lapse is then programmed for each microscope, with an identical workflow process. On the main touchscreen menu of the Juli, click on “Bright” and regulate the Power and Brightness values, which could be set up at about 30% and 10%, respectively.
7. Click on “Movie” to open a submenu with the following parameters to be set up: “Total time” (duration of the time lapse), “Interval” (gap between each acquired image), “Name” (name of the acquired image, saved as JPEG file and seuqnetially numbered by the system; to be entered by a submenu). There is a field named “# of images”, which is automatically updated to the total number of images to be acquired. In our case we need to generate a 24h time lapse (“Total time” = 24 h), with images acquired every 3 min (“Interval” = 3 min) (see *Note 31*).
8. When done click on “Bright” to indicate that only bright field is needed
9. Click on “Apply”. This will redirect to the main menu, where a button “REC” is now activated.
10. Click on “REC” to start the time lapse program and close the incubator inner and outer doors. Now the Juli video microscopes are programmed to save an images of the two co-cultures every 3 min until the time lapse will end at the 24 h time point (see *Note 32*).

### 3.10 Generation of the video from acquired images

1. Turn off the two Juli microscopes and transfer the images from the two SD cards on a PC (see *Note 33*).
2. To convert the sequential images to a frame-stacked video, open ImageJ and click on File > Import > Image sequence.
3. Choose the directory containing the image frames acquired by the Juli time lapses.
4. Set the values for “Start”, “Count”, “Step” and “Scale” to 1, 480, 1, 100, respectively.
5. Select “Sort names numerically” and “Use virtual stack”.
6. Click on “OK” to confirm data entered.
7. Now a video containing all the 480 frames appears, that need to be saved in the Audio Video Interleave (AVI) format. To do so click on File > Save as > AVI.
8. In the opened window choose “JPEG” in the “Compression” field and insert 5 frames per second (fps) as frame rate value for both videos (see *Note 34*).
9. Click on “OK” to save the video as AVI. In our case, we generated two 24h videos for hEo5 and hEo33 co-culteres, respectively.
10. In the ImageJ environment, open the desired AVI video.
11. To generate an “Inverteted” video, click on Process > batch > Macro.
12. Choose the “Invert” command to create a video where each white pixel in each frame of the video is converted to its inverse value (see *Note 35*).
13. Save this video and use it for cell tracking analysis.
14. Starting from this video, generate a region of interest (ROI) cropped video (300x300 pixels sized), which will be used for cell track detection. This video should be selected so that a centered A375P (target) cell appears. This can be done by the command Image > Crop in the ImageJ environment (see *Note 36*).
15. At the occurrence, optimize the brightness and contrast of the video file previously generated by clicking on Image > Adjust > Brightness/Contrast. Extend these setting to all the frames in the stack upon ImageJ request.

### 3.11 Cell tracking analysis and profiling of Eos5 and Eos33 from co-culture videos

1. Activate the TrackMate plugin (Plugins > Tracking > TrackMate).
2. During the first phase (spot selection) a first set of sequential menus opens, which allow to select eosinophils (spots) throughout all the image frames (see *Note 37*). Insert the required data as indicated in the pipeline (Figure 3).

**Figure 3.**
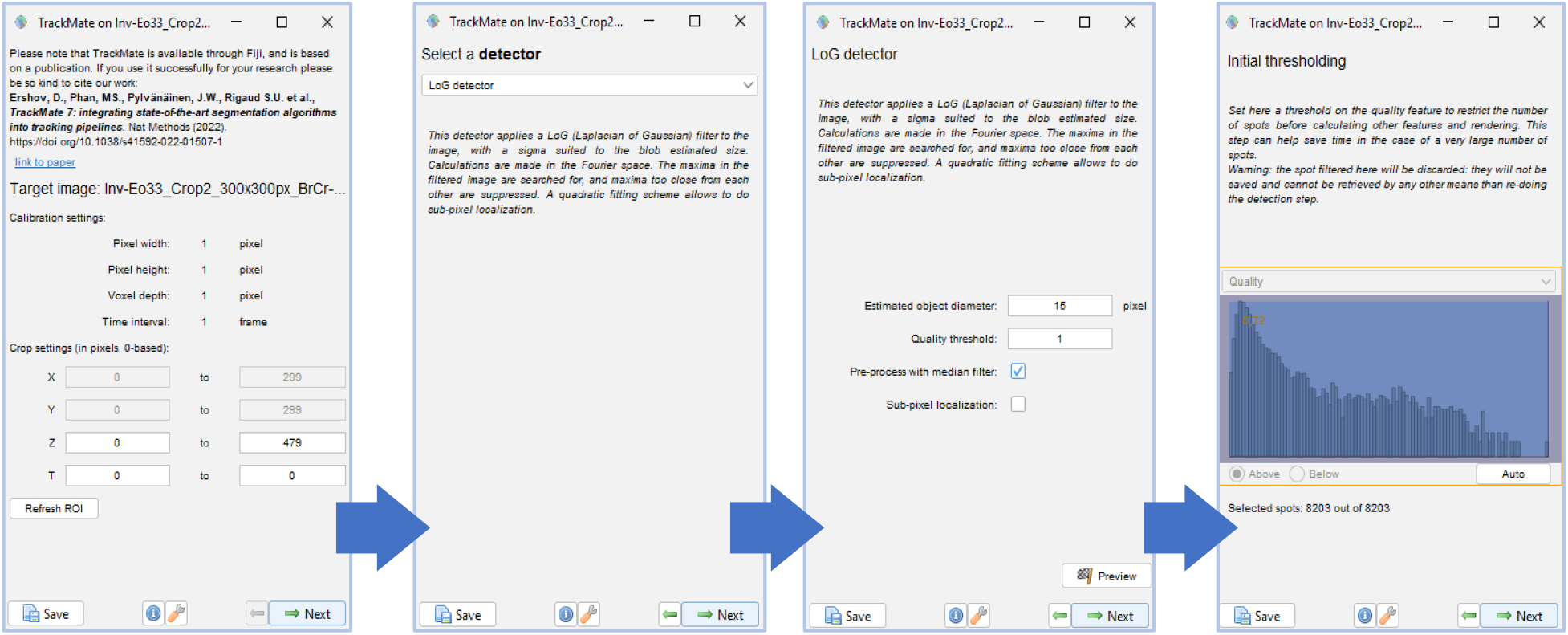
Example of the Spot (eosinophil) selection pipeline with the Laplacian of Gaussian (LoG) detector. These sequential menus allow the operator to select an appropriate spot selector algorithm. Blue arrows delineate the steps between the previous and the following menu. White boxes indicate the parameters to be set up.
3. At the completion of spot selection pipeline, a second set of menus opens, which serves to select the tracking algorithm (see *Note 38*) and subsequently to set it with the appropriate parameters (Figure 4).

**Figure 4.**
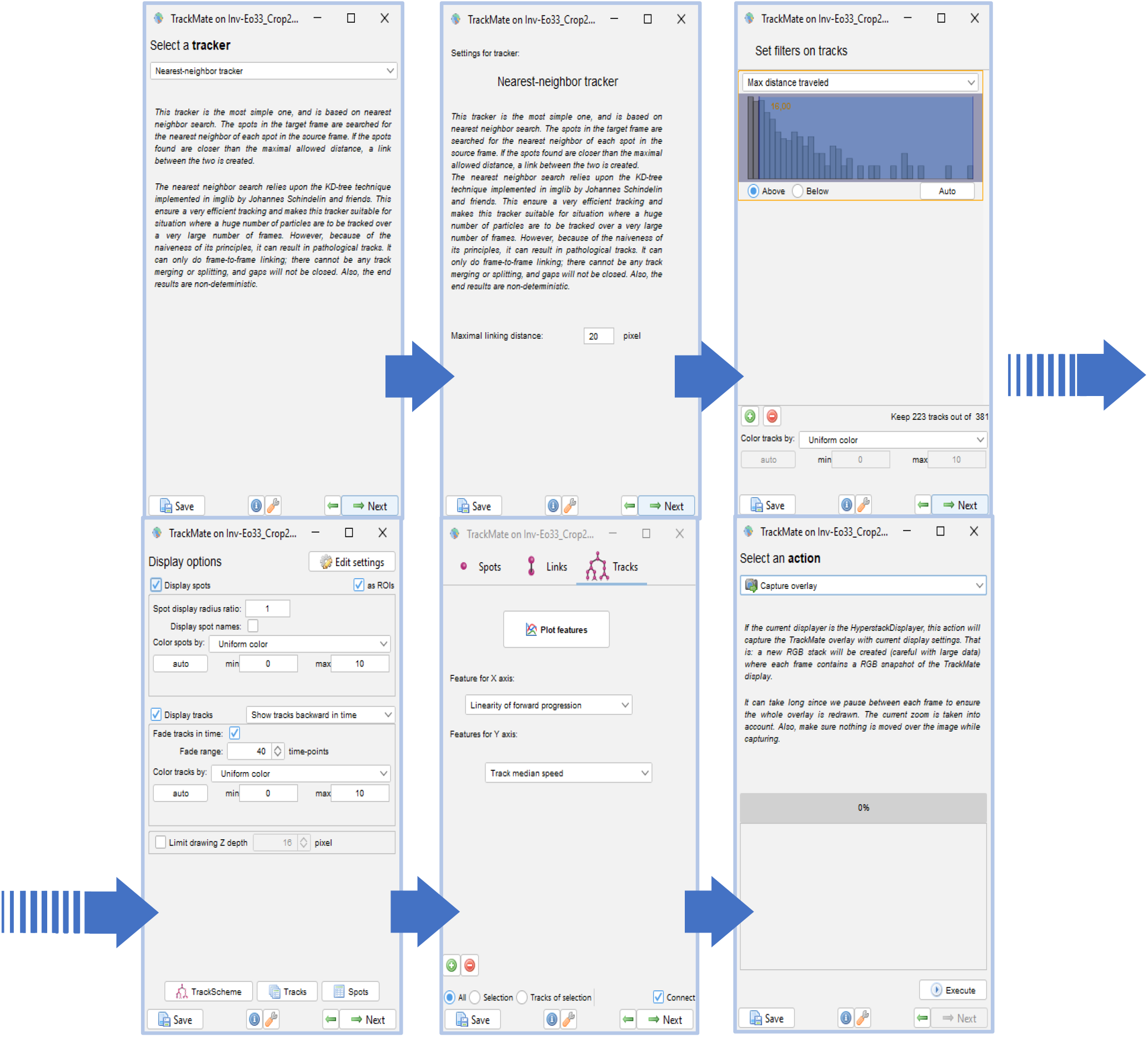
Example of Tracker selection pipeline with the Nearest-neighbor tracker. These sequential menus allow the operator to select an appropriate spot selector algorithm. Blue arrows delineate the steps between the previous and the following menu. White boxes show the parameters to be set up. Every window implements several buttons and options to further optimize the tracker efficiency in its ability to detect cell tracks.
4. At the end of the two pipelines a video with representative tracked eosinophils is available. Spot and track settings are then saved (see *Note 39*) and ready to be transferred in a visual graphics as overlays onto ROI microphotographs (Figure 5).

**Figure 5.**
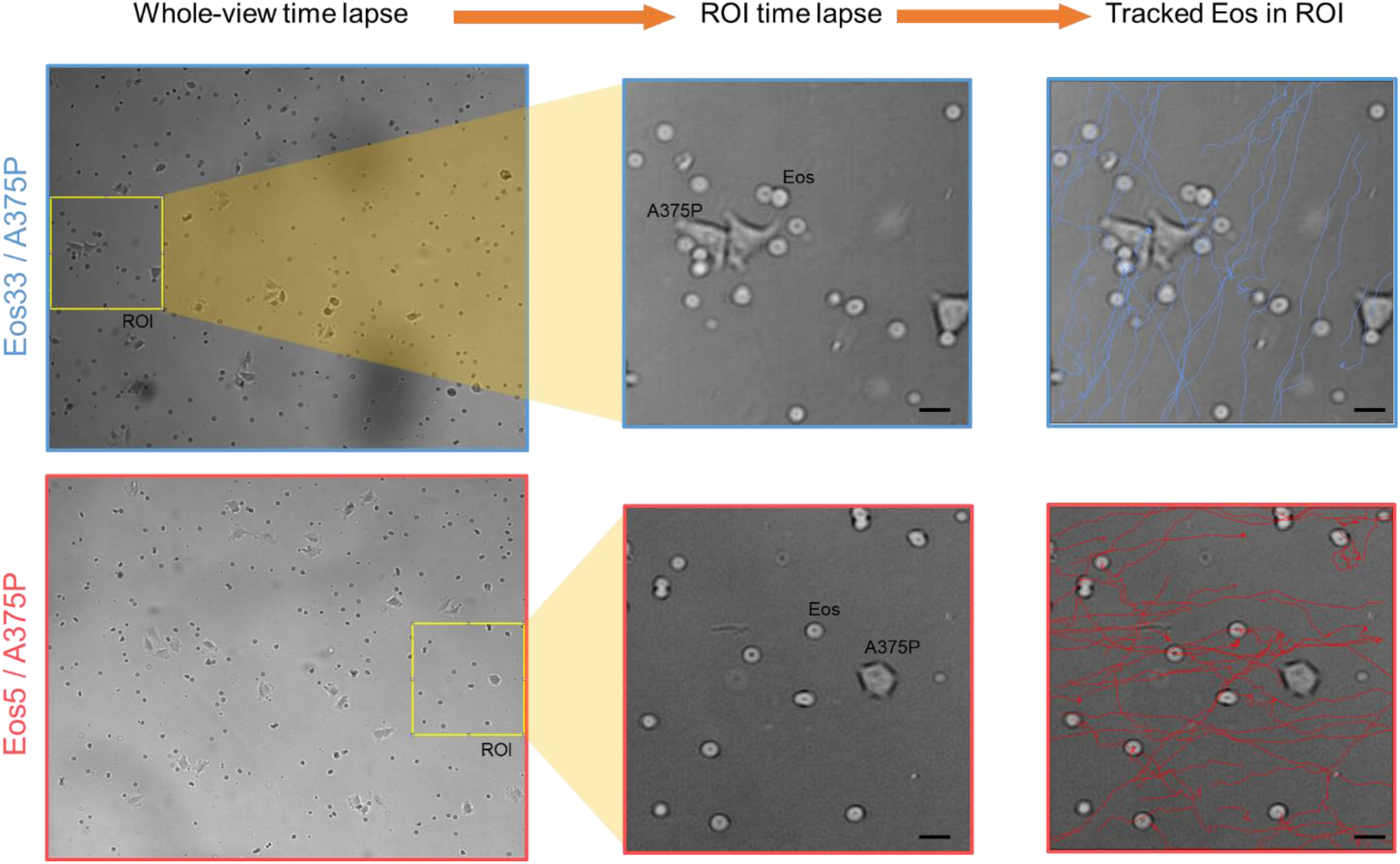
Example of automated Eos track profiling following co-culture with melanoma cells. A ROI video (ROI time lapse, 300x300 pixel) is extracted by the whole-view time lapse (1280x1024 pixel) and is subjected to Automated cell tracking by TrackMate plugin into the ImageJ environment (Tracked Eos in ROI). Scalebar, 20 μm.

### 3.12 Manual Calculation of interaction times between eosinophils and melanoma cells

1. Open the time lapse video file derived from Eos/A375P co-culture (Eos33/A375P or Eos5/A375P).
2. Visually select an Eos and keep note of the frame inside the time lapse video and the X-Y coordinates of that eosinophil for future reference.
3. Visually identify and Eos and start to advance the video one frame at once, following the Eos movement.
4. When the Eos starts to touch the tumor cell save the frame to which this event occurs (F_T_) and continue to advance the frames one at time.
5. When the Eos begin to detach from the cancer target cell, save the previous frame (F_D_).
6. Calculate the interaction time (IT) by multiply the difference between F_D_ and F_T_ with the time lapse interval between each frame (in this case 3 min) (see *Note 40*).
7. Repeat steps 2-6 for another Eos until an adequate number of IT values is reached for each Eos33 and Eos5 co-culture (see *Note 41*).
8. Collect IT data and graph them with GraphPad as dot plot graph type (Figure 6A).

**Figure 6.**
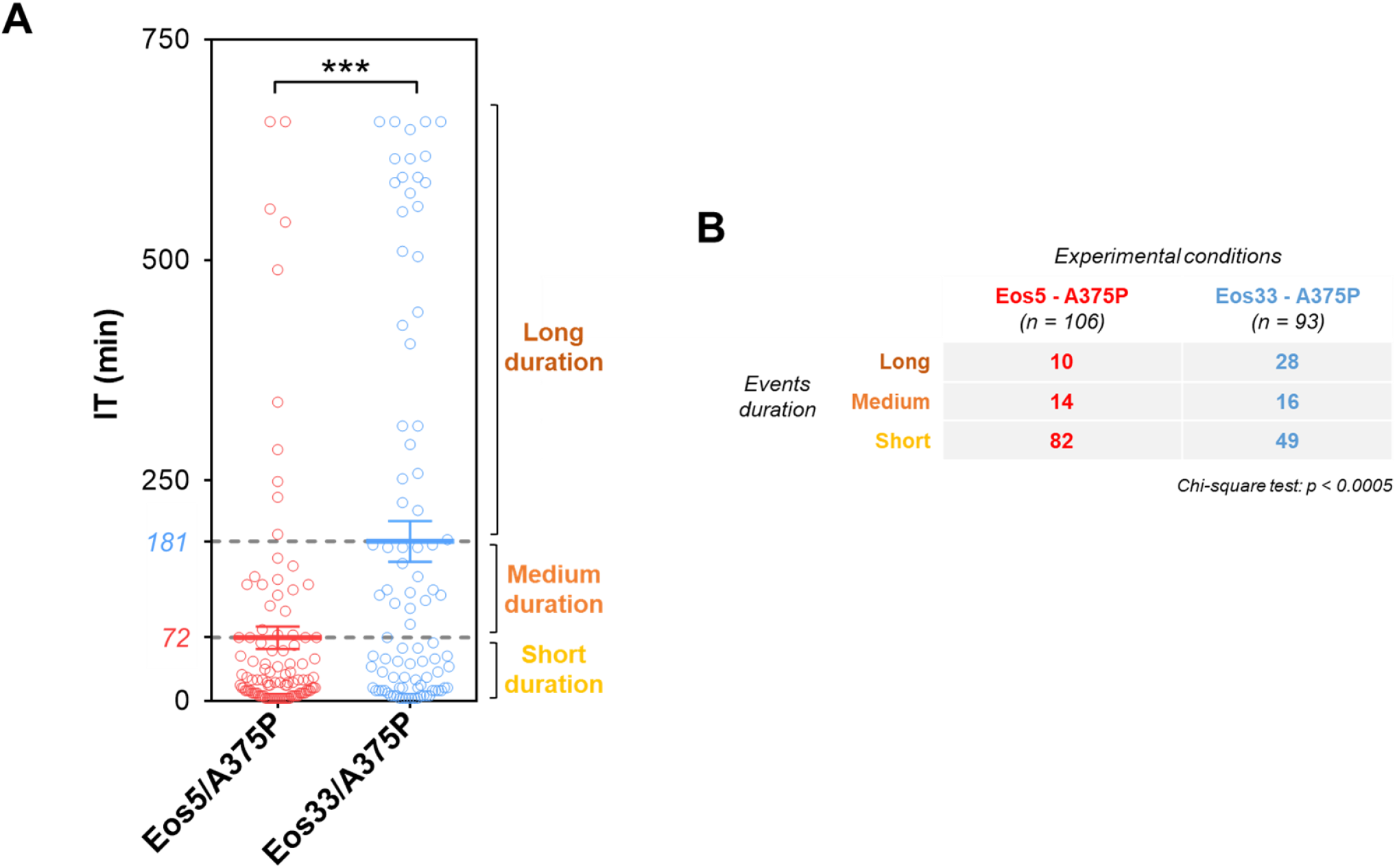
Example of IT values overview following co-culture of Eos with A375P melanoma cells. (A) Graph shows IT values manually extrapolated from the co-culture time lapse video. Gray lines and colored numbers depict the cut-off values extrapolated by the IT mean values of the two experimental groups and are used to subdivide the duration of contacts between tumor cells and Eos. ***, p < 0.0005, Mann-Whitney t test (dot plot graph). (B) Distribution of contact events over Eos5/A375P and Eos33/A375P experimental groups. The contingency table depicts the dispersion of IT events from panel A referred to the two indicated experimental conditions. The three duration event groups are delimited by the cut-offs deduced from panel A.
9. Manually generate two approximate cut-off values extrapolated by the mean IT values of the Eos5/A375P and Eos5/A375P conditions (Figure 6A) (see *Note 42*).
10. Check the significance p value between the two hEo33 and hEo5 groups exploiting the Mann-Whitney t test in GraphPad software.
11. Generate a contingency table (Figure 6B) and check the significance between the three contact groups by Chi-square analysis.

## 4. Notes

1. Note 1. To prepare Dextran/NaCl solution dissolve 0.45 g NaCl, 3 g Dextran in 50 mL 1X PBS and mix until the solution is clear and transparent. Store at 4°C overnight to allow Dextran microparticles to completely dissolve in the buffer. Dextran70/NaCl solution then is stable for at least 30-45 days if stored at 4°C. The day before use, filter the needed amount of solution with a 0.45 μm filter and store at 4°C. Bring to RT before use.
2. Note 2. ACK buffer is prepared by dissolving 8.26 g Ammonium Chloride, 1 g Potassium Bicarbonate and 0.037 g EDTA in 1 L of milliQ water. Once powders have completely dissolved, the solution is filtered in a 0.22 mm vacuum filter system. ACK is stable for at least 6 months if stored at 4 °C and under ordinary conditions of use. Store at 4°C for a maximum of 6 months.
3. Note 3. During this operation, avoid to form bubbles that can compromise the stratification process. On the other hand, it is fundamental to mix very well Dextran/NaCl with EDTA to a homogeneous solution in order to generate a correct fiber network. This will increase the efficiency in separation of red blood cells from leukocytes.
4. Note 4. Prepare a number of tubes with 10 mL PBS equivalent to the number of tubes for Dextran sedimentation. Use the same PBS-containing tube to transfer leukocytes gradually collected from each sedimentation tube, so that all leukocytes stratified from one tube are correspondingly transferred to a single tube. This is to avoid excess of residual Dextran, which may alter the subsequent centrifugation steps and, in turn, impair cell recovery.
5. Note 5. Supernatant must always be clear after this centrifugation. In case of turbidity, collect the supernatant, dilute 1:2 with PBS 1X, bring up to a final volume of 40 mL and repeat the centrifugation at 1500 rpm for 6 min at RT. Then, collect the pellet of residual cells.
6. Note 6. This passage must be performed with gentle manipulation of the tube, before, during and after the centrifugation. If needed, carefully use a serological pipette to discard the supernatant.
7. Note 7. Supernatant must always be clear after this centrifugation. In case of turbidity dilute the supernatant 1:2 with PBS 1X, bring up to a final volume of 40 mL and repeat the centrifugation at 1200 rpm for 8 min at RT
8. Note 8. This transfer must be done with extreme care. To correctly proceed with this transfer, resuspend the leukocytes pellet in exactly 5 mL of PBS 1X and then incline the 15 mL tube (already filled with the 5 mL Lymphosep). Place the pipette tip containing the cells on the top of the 5 mL Lymphosep and carefully flush down (with a slow flux rate) avoiding mixing. At the end of this operation, the Lymphosep phase and the leukocyte phase must appear separated.
9. Note 9. This centrifugation setting requires maximum acceleration, minimum deceleration and brake inactivation.
10. Note 10. At the end of the stratification process different phases are detectable. From top to bottom of the 15 mL tube, the stratification should present: a clear phase, a white and dense phase representing the Peripheral Blood Mononuclear Cells (PBMC), another clear and transparent phase with excess of Lymphosep solution or PBS and a final red pellet of granulocytes and erythrocytes. Separately collect each phase using 100 μl Gilson micropipette or serological pipette and discard.
11. Note 11. Use the maximum attention during this step, as the pellet could float during the elimination of the supernatant.
12. Note 12. Incubation times depend on the amount of ACK buffer. For 5 mL volume of this solution an 5 min incubation at RT is needed. It is recommended to use an amount of 4-6 mL of ACK.
13. Note 13. After blood cell lysis granulocytes are very sensitive. To limit activation and degranulation, work on ice from this step onward.
14. Note 14. The detection of a white pellet at the end of this centrifugation step denotes successful lysis of erythrocytes. Residual red blood cells may not be a problem. However, in case of substantial presence of red cells in the pellet, repeat steps 18-20.
15. Note 15. Dilute the 0.4% Trypan Blue solution 1:8 in PBS 1X. Use this solution to count cells appropriately diluted.
16. Note 16. This kit allows for negative selection of eosinophils. All steps involving the use of the kit must be carried out at 4°C, unless otherwise indicated.
17. Note 17. The number of Eos to be seeded depends on the yield obtained after magnetic separation. Normally from 1 buffy coat, 2-8 × 10^6^ eosinophils are obtained, but the number may vary from donor to donor.
18. Note 18. To prepare the working solution, add 2 μL of “x PKH26 Red fluorescent Cell Linker Solution in 498 μL PBS 1X in a 1.5 mL Eppendorf vial. Always keep this solution in the dark, i.e. by covering the vial with an aluminum fold.
19. Note 19. In this incubation step, shake the Eppendorf tube occasionally, paying attention to maintain the vial in the dark.
20. Note 20. If possible, use polypropylene tubes to avoid A375P cells adherence to the plastic support during this incubation. However, if using polystyrene tubes, immediately transfer the tubes on ice after the incubation and work on ice for the subsequent steps to facilitate the detachment of melanoma cells from the plastic support.
21. Note 21. From this step onward work on ice to avoid melanoma cells to attach to plastic support.
22. Note 22. Do not use EDTA-containing buffers, which can disaggregate conjugates.
23. Note 23. Do not mix with the micropipette tip since this may cause conjugate disruption. Simply mix the tube by gentle tapping.
24. Note 24. Attune CytPix Acoustic Focused Flow Cytometer (Gruijs et al., 2022) employs a combination of ultrasonic waves, akin to those utilized in medical imaging, and hydrodynamic forces to precisely arrange cells into a singular, concentrated line along the central axis. By facilitating the tight focusing of cells during laser examination, the system enables photon collection, thereby guaranteeing data quality regardless of the sample-to-sheath ratio.
25. Note 25. Single eosinophils are identified as PKH26^-^Siglec8^+^ from the “morpho” gate.
26. Note 26. Whit this tool it is possible to extrapolate and count multiple eosinophils (> 1) interacting with a single tumor cell, as a parameter of increased activation/adhesion and tumor reactivity.
27. Note 27. The two Juli video microscopes (Lee et al., 2021) represent the two co-culture experimental conditions, namely the Eos33/A375P condition and the Eos5/A375P control condition. Placing the Juli stations inside the incubator at least 24 hours before starting the experiment allows the microscope reaching the internal temperature of the incubator (37°C). This will minimize the formation of water condensation in the culture flasks, which can veil the images acquired during time lapse recording.
28. Note 28. Collect Eos5 and Eos33 by recovering all the medium and centrifuge at 1600 rpm for 5 min at 4°C. Resuspend the two eosinophil populations in RPMI co-culture medium at 3 × 10^5^ in 500 μL and dispense 500 μL per flask.
29. Note 29. The 3:1 (Eos:A375P) ratio is employed in this setting where A375P cells are adherent to the flask in order to facilitate the identification of eosinophils by the TrackMate in the tracking analysis of videos.
30. Note 30. We used a common 8 GB Secure Digital (SD) card for each Juli video microscope. Be sure the card is empty before its use.
31. Note 31. For a 24 h time lapse video recording, the “# of images” field will be automatically updated to 480 frames (namely the acquired images belonging to the video). Indeed, a 24 h interval represent 1440 min. Given that the interval between frames is 3 min, the total number of images is 1440 min / 3 min = 480 images in JPEG format.
32. Note 32. The time interval must be carefully chosen in function of the internal incubator humidity and of the exact type of support to be used. In our specific case, a 25cm2 flask is optimal for a 3 min interval of acquisition between frames. Prolonging this interval will reduce the increase of humidity in the flasks, but also does compromise the correct generation of cell tracks during cell tracking analysis by ImageJ. On the other hand, decreasing the interval will increase ImageJ ability to correctly generate cell tracks but also does increase the humidity inside the flasks, with possibility that water drops may form which can obscure the microscope objective.
33. Note 33. The PC can be equipped with Linux, Windows or MacOS operating systems, all supporting the Java language to be installed separately (Schmid et al., 2010). ImageJ is indeed a multiplatform software compatible with all these operating systems.
34. Note 34. Always use the same frame rate settings for all the videos belonging to an experiment. This will allow to perform comparison analyses between the two videos.
35. Note 35. The inversion process for the video is an essential requirement for a correct detection of the spots during the cell tracking analysis by ImageJ. Indeed, spots will be recognised as brilliant circles rather than original dark ones.
36. Note 36. It is strongly recommended to do tracking analysis on different ROIs of the video rather than on the overall video. This will allow to obtain cleaner tracks which can be easily viewable when they are in overlay to the video. Furthermore, performing a cell tracking analysis on an overall video may slow the operating system, especially when the PC is equipped with a low Random Access Memory (RAM) capacity.
37. Note 37. The available spot detectors algorithms are Laplacian of Gaussian (LoG) detector, Difference of Gaussian (DoG) detector, Hessian detector, Label Image detector, Mask detector, Thresholding detector, as well as manual selection of spots (eosinophils), without use of detectors. Quality (Pixel resolution and brightness/contrast ratio) of the images will be the determinant factor to choose the best spot detector.
38. Note 38. The available cell tracker algorithms are Nearest-neighbor tracker, linear assignment problem (LAP) tracker, simple LAP tracker, Kalman tracker, advanced Kalman tracker, Overlap tracker, as well as manual selection of eosinophil tracks.
39. Note 39. An eXtensible Markup Language (XML) file can be saved to store all the experimental settings. This will allow to store all the spot and track settings, which can be loaded by TrackMate (Fazeli et al., 2020; Rollins et al., 2022) to directly view the cell tracks without restarting from beginning.
40. Note 40. For example, if a Eos starts to touch the A375P at frame 234 and detachs at frame 302, the interaction time of that Eos (IT_Eos_) will be: IT_Eos_ = ( F_D_ – F_T_ ) * 3 min = ( 302 – 234 ) * 3 min = 204 min
41. Note 41. It is recommended to create a reference table representing a map to keep trace of the Eos used for calculation. This table could, for example, contain the X-Y coordinates of the Eos, the starting frames in which that Eos has been identified (namely the frame at which the Eos take contact with the target tumor cell), the end frame (namely the frame at which the Eos does detach from the target tumor cell).
42. Note 42. This will allow to categorize the IT values between the two experimental conditions into three groups: short, medium and long duration IT events. The medium IT value (71.68 min) of the control condition (Eos5/A375P) delimits the first group (short duration events from 0 to 72 min), whereas the second group (medium duration events from 72.5 to 181 min) is delimited by the mean of the control group and the Eos33/A375P condition. Lastly, the third group (long duration events from 181.5 min) contains all IT values above 181.5 min.

## 5. Concluding Remarks

Eosinophils are important components of the tumor microenvironment where they may play different roles in cancer progression (Varricchi, Galdiero, et al., 2018). Eosinophils can indirectly affect cancer progression through secretion of a variety of soluble mediators, including cytokines, chemokines and pro-angiogenic factors (Ghaffari & Rezaei, 2023; Varricchi, Galdiero, et al., 2018). On the other hand, eosinophils can directly kill cancer cells through adhesion-dependent degranulation (i.e., release of cytotoxic cationic proteins and granzymes) (Andreone et al., 2020; Ghaffari & Rezaei, 2023). Tumor cytotoxic function of eosinophils can be greatly enhanced upon activation with IL-33 (Andreone et al., 2019; Lucarini et al., 2017), IFN-γ (Reichman et al., 2019) or CCL11 (Xing et al., 2016). Evidence suggests that the effector function of eosinophils is contact-dependent (Andreone et al., 2019; Gatault et al., 2015) and thus strictly correlated with their ability to directly interact with cancer cells. In this respect, the interaction ability of eosinophils with cancer cells may represent a useful parameter for evaluating the activation status and effector propensity of these cells.

Our dual approach provides an accessible methodology to investigate the capability of eosinophils to functionally interact with cancer cells upon exposure to different stimuli (i.e., IL-5 or IL-33). On the one hand, acoustic focusing flow cytometry analysis allows to accurately select and count eosinophil/tumor cell conjugate formation by image visualization of flow cytometry gated populations discarding artifacts. On the other hand, time-lapse video microscopy allows to quantify eosinophil/tumor cell interactions by tracking profiles of these cells. These techniques, here employed using human eosinophils and A375P melanoma cells, can be easily readapted to other immune cell subsets that interact directly with cancer cells as well as to other tumor models, provided that the cancer cells grow in adherence to the culture moiety.

## Acknowledgements

This work was supported by AIRC (Associazione Italiana Ricerca Cancro) IG 21366 to GS and in part by the Innovation Ecosystem Heal Italia PE00000019, funded by the European Union - Next Generation EU, PNRR Mission 4 Component 2 Investment 1.3.

